# The role of integrins in *Drosophila* egg chamber morphogenesis

**DOI:** 10.1101/706069

**Authors:** Holly E. Lovegrove, Dan T. Bergstralh, Daniel St Johnston

## Abstract

A *Drosophila* egg chamber is comprised of a germline cyst surrounded by a tightly-organised epithelial monolayer, the follicular epithelium (FE). Loss of integrin function from the FE disrupts epithelial organisation at egg chamber termini, but the cause of this phenotype remains unclear. Here we show that the β-integrin Myospheroid (Mys) is only required during early oogenesis when the pre-follicle cells form the FE. *mys* mutants disrupt both the formation of a monolayered epithelium at egg chamber termini and the morphogenesis of the stalk between adjacent egg chambers, which develops through the intercalation of two rows of cells into a single-cell wide stalk. Secondary epithelia, like the FE, have been proposed to require adhesion to the basement membrane to polarise. However, Mys is not required for pre-follicle cell polarisation, as both follicle and stalk cells localise polarity factors correctly, despite being mispositioned. Instead, loss of integrins causes pre-follicle cells to basally constrict, detach from the basement membrane and become internalised. Thus, integrin function is dispensable for pre-follicle cell polarity but is required to maintain cellular organisation and cell shape during morphogenesis.

## Introduction

Epithelial cells are the most common animal cell-type, adhering to each other to form the sheets and tubes that comprise many tissues and organs. They therefore play essential roles as barriers between compartments and in the regulation of the directional traffic of molecules from one side of epithelium to the other. In order to carry out these functions, all epithelial cells must be polarised, with their apical domains facing the outside or lumen and their basal domains contacting the basement membrane (Rodriguez-Boulan and Macara, 2014). The orientation of the polarity axis depends on a lateral cue from cell-cell adhesion and external cues from the apical and/or basal sides of the epithelium. In well-characterised vertebrate epithelia, polarity is oriented by integrin-mediated adhesion to the basement membrane, and disruption of integrins leads to an inverted polarity and multi-layering (Chen and Krasnow, 2012; Eaton and Simons, 1995; Yu et al., 2005) (Akhtar and Streuli, 2012). On the other hand, primary epithelia in *Drosophila*, which derive from the cellular blastoderm, do not require integrins or a basal cue for their polarity and polarise in response to apical signals(Schmidt and Grosshans, 2018). Some *Drosophila* epithelia form later in development from mesenchymal to epithelial transitions, and it has been proposed these secondary epithelia require a basal cue to polarize (Tepass, 1997). In support of this view, the endodermal cells of the embryonic midgut must contact the basement membrane of the visceral mesoderm to polarize, and the enterocytes of the adult midgut require components of the integrin adhesion complex to integrate into the epithelium and polarise (Chen et al., 2018; Tepass and Hartenstein, 1994).

It is less clear whether integrin adhesion to the basement membrane is required in the other well-characterised secondary epithelium in *Drosophila*, which is formed by the follicle cells that surround developing female germline cysts. All cells in a *Drosophila* egg chamber are generated in a structure known as the germarium, which resides at the anterior tip of each ovariole (Fig. 1A). The follicle stem cells (FSCs), which produce the somatic cells in each egg chamber, lie partway along the germarium (until this point the germline cysts are surrounded by escort cells, Fig. 1A)). FSC progeny migrate to surround each germline cyst as it moves through region 2 of the germarium. These progeny cells give rise to both the main follicle cells and, via a signalling relay, the polar cells and interfollicular stalk cells (Fig. 1A) (Grammont and Irvine, 2001; McGregor et al., 2002; Torres et al., 2003).

**Figure 1.**
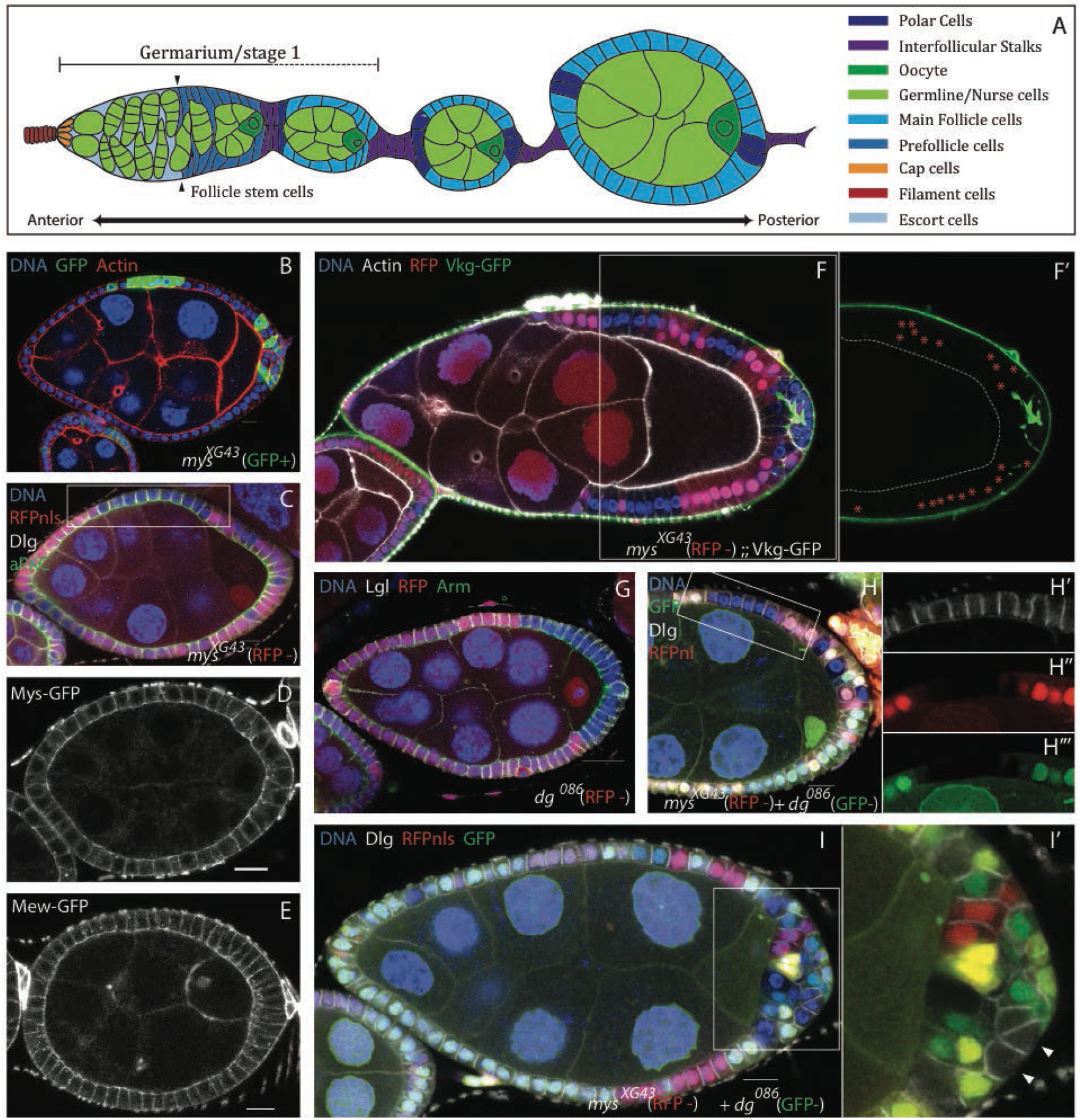
**(A)** Diagram showing a *Drosophila* ovariole, with the germarium on the left and successively older egg chambers on the right. The different cell types are indicated by colour (colour key; right). **(B)** A stage 7 egg chamber containing *mys*^*XG43*^ mutant cells (GFP+; green) stained for F-actin (red) and DNA (blue). The mutant cells produce disorganisation of the follicular epithelium (FCE) at the termini of egg chambers. (C) A stage 8 egg chamber containing lateral *mys*^*XG43*^ mutant cells (marked by the loss of RFP; red) stained for aPKC (green), Dlg (white) and DNA (blue). The mutant cells are organised and polarised correctly (box). **(D and E)** Stage 6 egg chambers expressing endogenously tagged Mys-GFP (C) and Mew-GFP (D). Both proteins show a uniform localisation around the plasma membrane of the follicle cells and are not enriched basally. **(F)** A stage 9 egg chamber containing *mys*^*XG43*^ mutant cells (RFP-), expressing Vkg-GFP (Collagen IV; green) from a protein trap insertion and stained for F-actin (white) and DNA (blue). The mutant cells, including those in the disordered region at the posterior do not secrete Vkg-GFP apically, but Collagen IV is secreted between the cell layers at the posterior. (F’) shows Viking-GFP alone. The dashed line marks the boundary between the oocyte and the follicle cells and the red stars mark the RFP+ wild-type cells. **(G)** A stage 8 egg chamber containing *dg*^*086*^ mutant cells (RFP -), stained for Lgl (white), Arm (green) and DNA (blue). The mutant cells do not disrupt the organisation of the FCE or apical-basal polarity when they occur at the egg chamber termini. **(H and I)** Dg does not act redundantly with Mys. Stage 8 egg chambers containing mutant clones of both *mys*^*XG43*^ (RFP -) and *dg*^*086*^ (GFP-), stained for Dlg (white) and DNA (blue). (H) Lateral double mutant clones (RFP and GFP negative) do not disrupt epithelial disorganisation or polarity (box). (I) Double mutant clones at the posterior cause epithelial disorganisation that is not discernibly worse than *mys*^*XG43*^ clones alone. Dlg is still excluded from the basal side of double mutant cells that contact the basement membrane (arrow heads in I’). All scale bars are 10μM.

Early work suggested that the polarity of the follicular epithelium depends on a basal cue provided by contact with the basement membrane, as well as lateral adhesion between cells and an apical cue provided by contact with the germline cysts (Tanentzapf et al., 2000). Consistent with this, mutations in *myospheroid* (*mys*), the only β integrin subunit expressed in the *Drosophila* ovary, cause disorganisation of the follicle cell epithelium (FCE) (Delon and Brown, 2009; Devenport and Brown, 2004; Fernández-Miñán et al., 2007). This disorganisation only occurs in mutant clones at the egg chamber termini, however, and lateral clones are indistinguishable from neighbouring wild type cells (Fig. 1B and box Fig. 1C).

Integrins establish a connection between the cytoskeleton and the extracellular matrix (ECM), which generally lies on only one side of a cell (Maartens and Brown, 2015). Integrin signaling could therefore potentially provide a symmetry-breaking polarity cue, and it has therefore been proposed that loss of integrin function in the terminal follicle cells leads to a loss of polarity either directly or through misoriented cell divisions (Fernández-Miñán et al., 2007; Fernández-Miñán et al., 2008). However, we observed that spindles are correctly oriented in *mys* mutant follicle cells (Bergstralh et al., 2013). Furthermore, spindle misoriention in the FCE does not cause disorganisation of the tissue, as cells that are born outside of the epithelium reintegrate (Bergstralh et al., 2015). These findings prompted us to re-evaluate the role of integrins during oogenesis to investigate the cause of the *mys* mutant phenotype.

## Results

### Myospheroid does not regulate apical-basal polarity

As previously reported, mutant clones homozygous for a null allele of *mys* cause disorganization of the epithelium in terminal regions (Fig. 1B) (Fernández-Miñán et al., 2008). This phenotype is also observed upon disruption of apical-basal polarity regulators (*e.g.* Discs large and Lethal giant larvae) (Bilder et al., 2000). We therefore considered the possibility that Myospherioid regulates follicle cell polarity. However, our evidence indicates that it does not:

Firstly, as previously reported, the loss of Mys function has no effect on the localization of polarity factors in main body follicle cells (Fig. 1C). Furthermore, the β-integrin Mys and the α-integrin Multiple edematous wings (Mew), unlike other polarity factors, do not localise in a polarised manner, but are instead spread evenly around follicle cell surfaces (Fig. 1D & E).

Disruption of Crag or Rab10, which promote the basally-directed secretion of the ECM, causes epithelial disorganization without obviously affecting the localisation of polarity determinants (Denef et al., 2008; Lerner et al., 2013). This finding raised the possibility that Mys, which binds the ECM directly, might also regulate polarised ECM secretion. We therefore examined the location of the ECM in *mys* mutant clones using a GFP protein trap in the *viking* locus (Vkg-GFP) (Collagen IV, α2 chain) (Morin et al., 2001). In the areas of disorganization, accumulations of Vkg are present within the epithelial layer, presumably because it is secreted by the disorganised cells and then trapped between them (Fig. 1F). However, unlike in *crag* mutant follicle cells, Vkg was not found apical to the *mys* mutant cells (i.e. between the follicle cells and the germline cells (dashed line Fig. 1F’)). This is true whether or not the mutant cells were part of a disorganised area. Thus, epithelial disorganisation in *mys* mutant clones cannot be attributed to mispolarised secretion of ECM components.

Integrins are not the only transmembrane proteins capable of interacting with the ECM. Dystroglycan (Dg), like the Mys/Mew integrin heterodimer, is expressed until stage 12 of oogenesis, binds to laminin and, via intracellular binding partners, links the ECM to the actin cytoskeleton (Ibraghimov-Beskrovnaya et al., 1992). It is therefore possible that Integrins and Dg act as redundant ECM receptors to provide a basal polarity cue. As previously reported, mitotic clones of the null allele *dg*^*086*^ affect neither the polarity nor the organisation of the follicle cells, regardless of whether they occur at the termini or along the sides of an egg chamber (Fig. 1G & Fig. S1A) (Christoforou et al., 2008; Haack et al., 2013). We therefore generated *mys* null mutant clones marked by the loss of RFP and *dg* null mutant clones marked by the loss of GFP in the same egg chambers, and examined the double mutant clones that lack both RFP and GFP (Fig. 1H & I). Cells mutant for both *mys* and *dg* still polarise correctly, provided that they are in contact with the germline and therefore can receive any apical polarity cues (Fig. 1H). Furthermore, the presence of doubly mutant cells at the termini of egg chambers does not make the epithelial disorganisation discernably worse (Fig. 1I). More importantly, the lateral polarity marker Discs large (Dlg) is still excluded from the basal side of *mys dg* double mutant cells in the disorganised regions that lack contact with the germline, indicating that the basal domain has been specified as distinct from lateral (arrow heads Fig.1I’). Thus, the ability of integrin mutant cells to polarise is not due to a redundant function with Dystroglycan. This suggests that the follicle cells do not need a basal cue to polarise and that the *mys* phenotype is caused by some other defect.

### The *mys* disorganization phenotype originates in the germarium

We next asked when the *mys* phenotype arises during oogenesis. To address this question, we depleted *mys* mRNA from the entire epithelium at different stages of development. GR1-GAL4 is expressed in the follicle cells of fully formed egg chambers, starting weakly at approximately stage 2/3 of oogenesis and getting stronger through development (Fig. 1B). Expressing an shRNA directed against *mys* under the control of GR1-GAL4 is lethal at 25 or 29°C. probably because GR1-GAL4 reduces Mys function in other tissues during larval or pupal development. However, adult flies eclose if the larvae and pupae are grown at 18°C, at which temperature the GAL4 system is inefficient. We therefore raised GR1-GAL4; UAS-*mys-*shRNA flies at 18°C, then shifted them to 29°C for 2 or 6 days before dissecting their ovaries. Under these conditions, Mys is strongly depleted from stage 3/4 of oogenesis onwards. (Fig. 2B). The morphology of the FCE is normal in these flies (Fig. 2C and Fig. S2A). Apical-basal polarity is likewise unaffected, as shown by the correct localisation of the apical marker aPKC and the lateral marker Lethal giant larvae (Fig. 2C and Fig. 2A).

**Figure 2.**
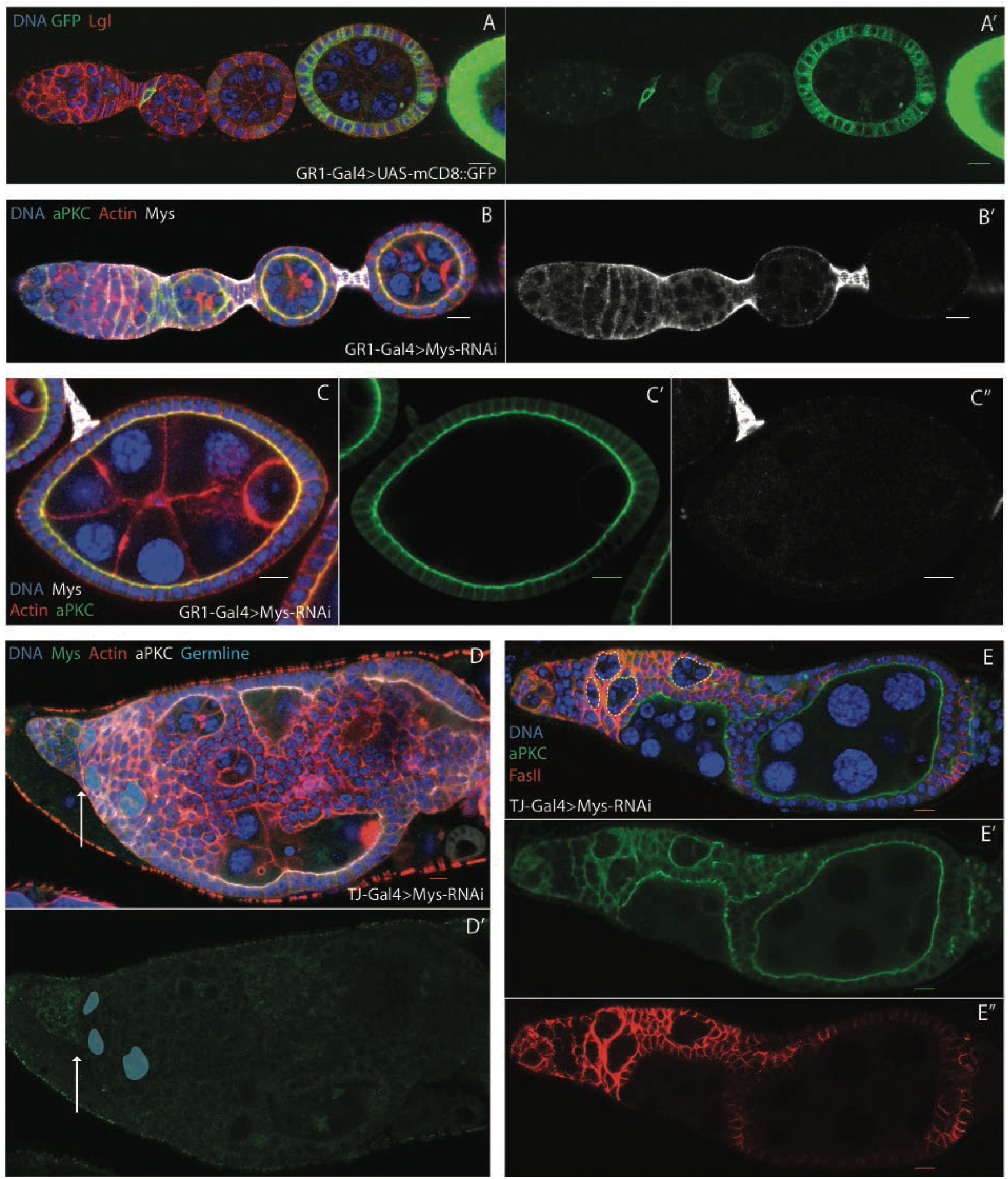
**(A)** The anterior portion of an ovariole showing the expression of UAS-mCD8::GFP (green) driven by GR1-Gal4, stained for Lgl (red) and DNA (blue). GR1-Gal4 drives weak expression in the follicle cells at stage 2/3 and increases in strength in later stages. It is expressed in the polar cells at all stages, but not in the interfollicular stalks. **(B and C)** An ovariole and egg chamber expressing UAS-*mys*-RNAi (Bloom 33647) under the control of GR1-Gal4, stained for Mys (white), aPKC (green), F-actin (red) and DNA (blue). The flies were raised until eclosion at 18°C and transferred to 29°C as adults for 2 days before dissection. Mys staining is lost from the follicle cells from stage 3/4 onwards (B’ and C”), but this has no effect on tissue organisation or apical-basal polarity (C’). **(D)** An ovariole expressing UAS-*mys*-RNAi under the control of TJ-Gal4, stained for Mys (green), aPKC (white), F-actin (red) and DNA (blue). Mys is strongly depleted from the somatic cells at all stages, resulting in gross disorganisation of the ovariole from the approximate position of the follicle stem cells (arrow). Blue shading marks the early germline cysts. **(E)** An ovariole expressing UAS-*mys*-RNAi under the control of TJ-Gal4, stained for aPKC (green), FasII (red) and DNA (blue). The follicle cells that contact the germline are correctly polarised. Early germline cysts are outlined in dashed white lines. All scale bars are 10μM.

We used Traffic Jam-GAL4 (TJ-GAL4) to investigate a role for Mys prior to stage 2/3. TJ-Gal4 is expressed in all somatic cells in the *Drosophila* ovary, apart from the interfollicular stalk cells (Fig. Arrow head S2B). Flies expressing *mys*-shRNA under the control of TJ-GAL4 demonstrate gross disorganisation of ovarioles, with multi-layering of the follicle cells and incomplete encapsulation and fusion of the germline cysts. (Fig. 2D & E). An identical phenotype is produced by a second *mys*-shRNA that targets a different region of the mRNA, confirming that this phenotype is due to Mys knock-down (Fig. S2C). TJ-GAL4 drives expression in the germarium in both pre-follicle cells and escort cells, but these latter cells are unlikely to be involved in the phenotype, since disorganisation first appears in the region where the FSCs reside (arrow Fig. 2D). We also considered the possibility that the phenotype arises during ovariole development, since TJ-GAL4 is expressed in the pupal ovary (Vlachos et al., 2015). This was tested by allowing the flies to develop and eclose at 18°C. These flies developed morphologically-wild type germaria (only 1 of 12 germaria examined displayed mild disorganisation) and egg chambers (Fig. S2D and E). When sibling flies were shifted to 29°C for 2 days prior to dissection, the majority of germaria examined (19 out of 20) displayed disorganisation and multi-layering (Fig. S2F & G). Later-stage egg chambers, which were produced before the temperature shift, appeared normal (Fig. S2F & G). This result shows that disorganisation develops in the adult germarium without any effect on the pupal ovary. Cumulatively, these findings indicate that the epithelial disorganisation in *mys* mutants arises in the germarium before stage 2/3 of oogenesis, and that this is the only stage when Mys function is required for the formation of a normal follicular epithelium.

Despite their gross disorganisation, TJ-GAL4; UAS-*mys*-shRNA egg chambers maintain certain features of a normal ovariole. Individual germline cysts still remain distinct from the somatic tissue and undergo their usual morphological changes, such as transforming from a disc of cells into a sphere (Fig. 2E and F). They are also capable of generating eggs, although these are generally misshapen, most probably because integrins are required for the egg chamber rotation that is required for normal egg elongation (Fig. S2H) (Lewellyn et al., 2013). Furthermore, those follicle cells contacting the germline are able to polarise, as evident by the correct localisation of aPKC and Fasciclin II (FasII) (Fig. 2E). Integrins are required to anchor the follicle stem cells (FSC) to their niche, and *mys* mutant FSCs move into the centre of the germarium and more frequently lost than wild-type FSC, or become quiescent (Hartman et al., 2015; O’Reilly et al., 2008). However, the FSCs still produce an approximately normal number of pre-follicle cells when Myospheroid is knocked down. This may be because TJ-GAL4 is not expressed in the FSC or because there is no competition between wild-type and mutant FSC, as Mys is knocked down in all cells. Thus, in addition to this role in FSC maintenance, Mys is required in the FSC and/or the pre-follicle cells for normal egg chamber morphogenesis.

### Mys participates in stalk formation

The observations that ovarioles fail to separate into discrete egg chambers when Mys is strongly depleted in the germarium (and onwards) and that epithelial disorganisation only occurs when mitotic clones occur at the egg chamber termini, raised the possibility that disorganisation reflects a defect in interfollicular stalk formation. We tested whether the interfollicular stalk cells are able to differentiate from the other pre-follicle cells, which go on to form the follicle cells and polar cells, by staining for the stalk cell marker Lamin C (Fig. 3A). This protein is still expressed in patches of cells between germline cysts when Mys is depleted using TJ-GAL4; UAS-*mys*-shRNA (Fig. 3B). Furthermore, Eyes absent (Eya), which is expressed by the follicle cells but not the interfollicular stalk and polar cells, is expressed in cells contacting the germline but not in patches of cells between the cysts (Fig. S2I and J). Thus, the somatic cells appear to be able to undergo their typical position-dependent differentiation programmes into follicle, polar, and interfollicular stalk cells in the absence of integrins. This suggests that the lack of interfollicular stalks upon Mys depletion is due to a failure in the cell shape changes and movements that usually generate this structure. Since little is known about the morphogenesis of interfollicular stalks, we set out to characterise this process in more detail.

**Figure 3.**
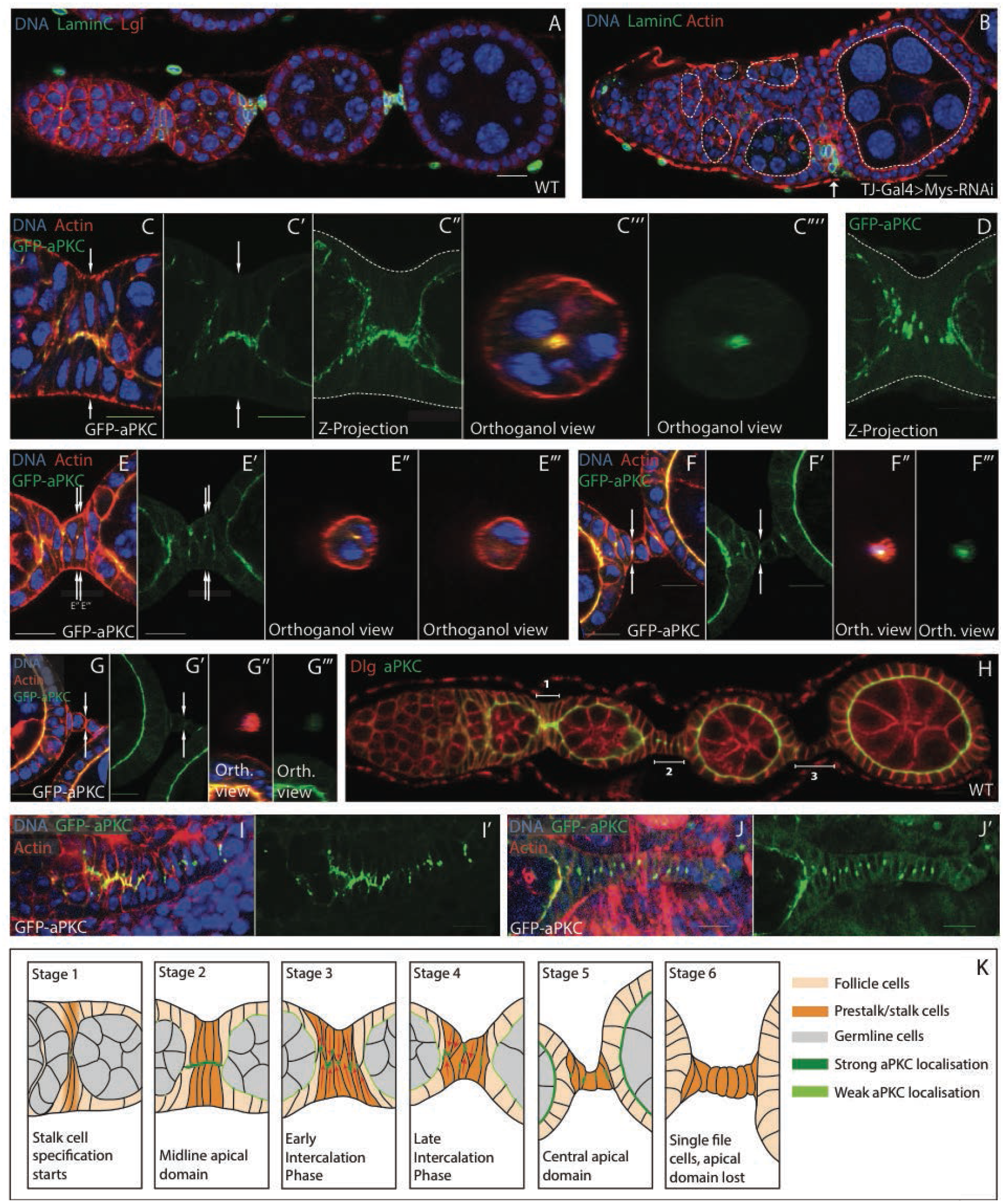
**(A)** A wild-type ovariole stained for Lamin C (green), Lgl (red) and DNA (blue). Lamin C is only expressed in the interfollicular stalks (A’). **(B)** An ovariole expressing UAS-*mys*-RNAi under the control of TJ-Gal4 at 25°C stained as in (A). Lamin C positive cells can be found between the developing germline cysts (arrow), showing that the interfollicular stalk cells are specified normally. Germline cysts outlined with dashed lines. **(C - G)** High magnification views of the steps in the morphogenesis of the stalk in wild-type ovarioles expressing endogenously-tagged GFP-aPKC and stained for F-actin (red) and DNA (blue). (C) Prior to intercalation, the pre-stalk cells form a two-cell wide column with aPKC-GFP localised to the points where opposing cells meet (C’; C” shows a Z-projection of 19 planes taken 0.5μM apart; C”’ and C”” show an orthogonal view through the region indicated by the white arrows in C and C”). (D) A Z-projection of 18 planes collected 0.5μM apart. As intercalation begins, GFP-aPKC localises to punctae that lie roughly along the midline of the tissue. Dashed lines in C” and D mark the basal sides of the cells. (E-G) The localisation of aPKC-GFP in interfollicular stalks of increasing age. E’’, E’’’, F’’, F’’’, G’’ & G’’’ are orthogonal views of Z stack images taken 0.5μM apart. Arrows indicate the position of each orthogonal view. In E the left arrow shows the position of E’’ and the right E’’’. **(H)** An ovariole stained for Dlg (red) and aPKC (green) showing the stages of stalk formation. Dlg localises to the cortex of interfollicular stalk cells where they contact their neighbours in forming (brackets 1 and 2) and mature stalks (bracket 3). **(I)** An early developing pupal ovary expressing aPKC-GFP (green, I”) and stained for actin (red) and DNA (blue). aPKC localises to the apical sides of the basal stalk cells prior to intercalation. Full image in Fig. 3F **(J)** An older pupal ovary expressing aPKC-GFP (green) and stained for F-actin (red) and DNA (blue)., aPKC relocalises to lateral punctae as the basal stalk cells intercalate. Full image in Fig. 3G **(K)** Diagram showing the positions of cells in examples of each stage of interfollicular stalk formation All scale bars are 10μM.

The stalk begins as a double row of elongated cells between developing germline cysts that arrange themselves so that their apical sides meet along the midline of the germarium (Fig. S3A). As previously described, these cells then undergo an intercalation, such that the two-cell thick column becomes a single row of cells (Godt & Laski 1995) (Fig. S3A). In the early stages of stalk formation, aPKC localises to the apical tips of the stalk cells, facing the centre of the column (Fig. 3C and Fig. 3H; bracket 1). As intercalation proceeds, aPKC does not remain associated with the tips and instead forms small dots on the lateral membrane that lie along the centre of the column (Fig. 3D). The intercalating cells eventually resolve into a single line of cells with staggered nuclei. At this point, the aPKC foci have moved to the middle of the lateral interface between adjacent cells (Fig. 3E & Fig. 3H bracket 2). Finally, the cells fully intercalate such that their nuclei are arranged in a line (Fig. 3F). A small focus of aPKC persists at the centre of the interface between cells, but this eventually disappears and aPKC is almost undetectable (Fig. 3F and 3G). This leaves only lateral polarity factors, such as Dlg, localised along the length of the stalk cell-cell contacts (Fig. 4I; bracket 3). By contrast, lateral cell adhesion components such as FasII and Fasciclin III (Fas3), which are expressed along the lengths of cell-cell contacts while the cells are in the germarium, are lost over the course of intercalation (Fig. S3B and C).

**Figure 4.**
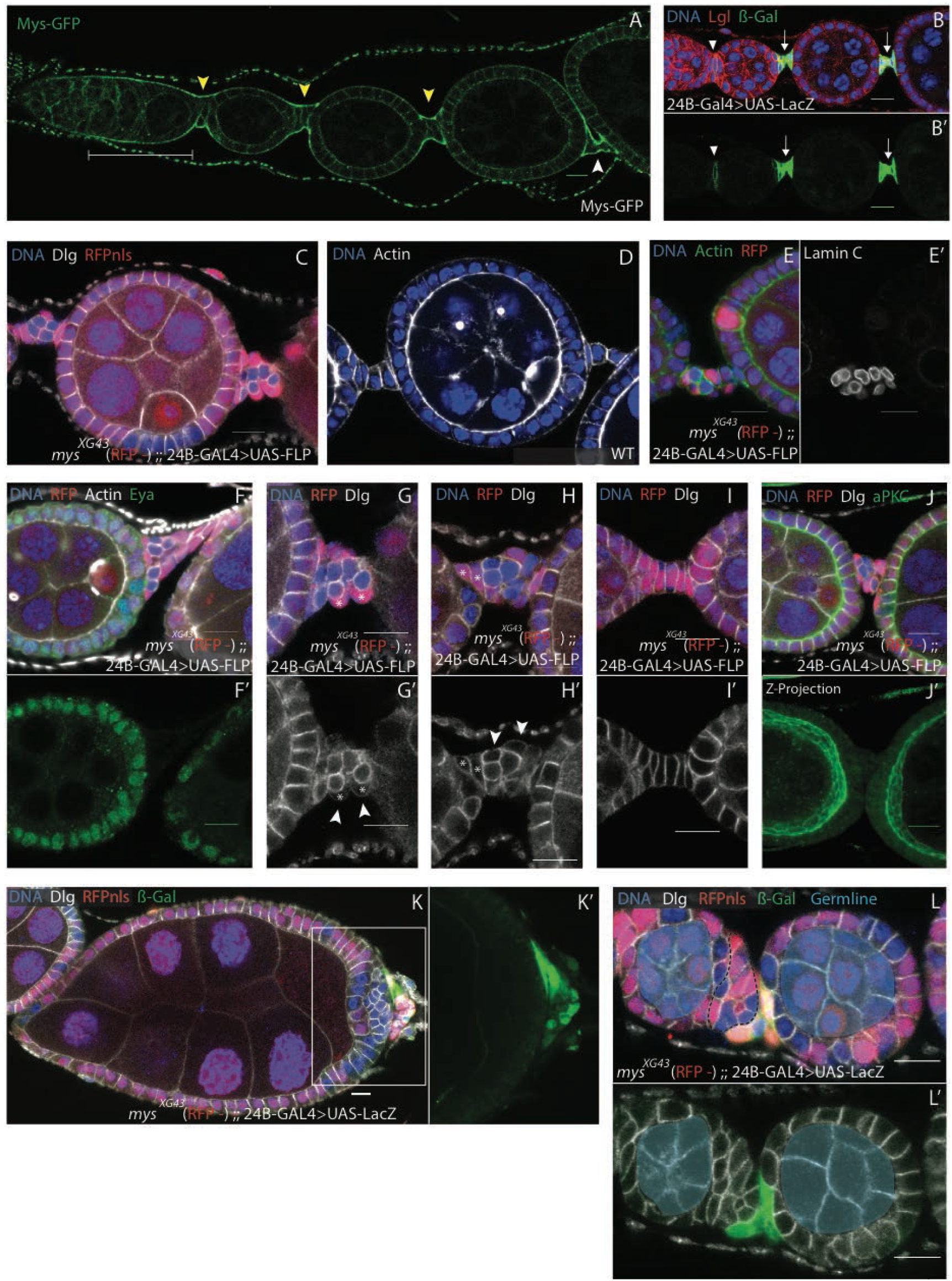
**(A)** A wild-type ovariole expressing Mys-GFP (green) from a genomic BAC transgene. Mys is enriched on the basal side of all pre-follicle cells in the germarium (bracket) and in the forming and mature interfollicular stalks (yellow arrowheads and white arrowhead, respectively) **(B)** A section of ovariole stained for Lgl (red), DNA (blue) and β-galactosidase (green and B’) showing the pattern UAS-LacZ expression under the control of the 24B-Gal4 driver. 24BGal4 drives expression in the interfollicular stalk cells only (arrow head and arrows). **(C)** A stage 4 egg chamber stained for Dlg (white) and DNA (blue) containing *mys*^*XG43*^ clones marked by the loss of RFP (red). 24B-Gal4 was used to drive expression of UAS-FLP to generate *mys*^*XG43*^ clones specifically in the interfollicular stalks. The presence of *mys* mutant cells disrupts the organisation of the interfollicular stalks. **(D)** A wild-type stage 4 egg chamber stained for F-actin (white) and DNA (blue) showing the full-formed stalks that separate it from the adjacent younger and older egg chambers. **(E-J)** *mys*^*XG43*^ clones (make by the lack of RFP (red)) in interfollicular stalks induced by the expression UAS-FLP under the control of 24B-Gal4. (E & F) A disorganised stalk containing *mys* clones stained for F-actin (green), DNA (blue) and Lamin C (white in E’). Lamin C is expressed in both mutant and wild type cells in disrupted interfollicular stalks (E). (F) A disorganised stalk containing *mys* clones stained for Eya (green) and DNA (blue). Eya is turned off normally in both the mutant and wild-type stalk cells. (G and H) Mutant interfollicular stalk cells are round in appearance compared to a stalk made from all wild type cells (I). Wild type cells in disrupted interfollicular stalks range from rounded (white stars in G) to wild type (white stars H). Dlg only localises to the regions where neighbouring interfollicular stalk cells contact each other in disrupted interfollicular stalks (arrow heads G’ and H’) as in interfollicular stalks containing only wildtype cells (I’). (J) A disorganised interfollicular stalk stained with aPKC (green) and Dlg (white), (J’) a Z-projection of 16 planes taken 1μM apart of the stalk in (J). This shows that aPKC is not present in mature stalks containing *mys*^*XG43*^ mutant cells. **(K and L)** Regions of ovarioles stained for DNA (blue) and β-galactosidase (green) containing hs-FLP-induced *mys*^*XG43*^ clones marked by the loss of RFP (red). The interfollicular stalk cells are marked by β-galactosidase (green) expressed from UAS-LacZ under the control of 24BGal4. The disorganised region caused by *mys*^*XG43*^ clones at the terminus of the stage 9 egg chamber in (Box K) does not contain cells expressing the β-galactosidase stalk marker, which lie only in a malformed stalk posteriorly (K’). (L) The stalk cells lie at the posterior of the region between the two younger egg chambers (LacZ positive cells, green). The black dashed line identifies cells that could potentially contribute to the disorganised region at the terminus of the egg chamber later in development, like that seen in (K). All scale bars are 10μM.

Adherens junctions components, such as E-cadherin, demonstrate a similar pattern of localisation to aPKC (Fig. S3D and E). A similar dynamic localisation of E-cadherin has also been described in the developing basal stalks in the *Drosophila* pupal ovary, which link developing ovarioles to the posterior end of the ovary (Vlachos et al., 2015) (King, 1970). Like intrafollicular stalks, basal stalks are formed by a cell intercalation event that generates a single row of cells from (in this case) a disorganised area of cells (Godt and Laski, 1995). Examination of developing basal stalks reveals that aPKC also localises to the central tips of these cells as they line up two abreast prior to intercalation (Fig. 3I and S3F). Once a single row of cells has been established, single accumulations of aPKC can be seen between each cell along the midline of the basal stalk (Fig. 3J and S3G). These findings suggest that the intercalation events in the pupal basal stalks and adult interfollicular stalks are driven by a highly similar sequence of morphological events.

As outlined above, interfollicular stalks are made by a complex series of cell movements and shape changes (Fig. 3K). Throughout this process, integrins (both Mys and the alpha subunit Mew) are strongly enriched along the basal sides of the stalk cells, in contrast to their uniform distribution around the membrane later in development (Fig. 4A and Fig. S4A). To determine whether Mys is required cell-autonomously in the presumptive stalk cells for stalk morphogenesis, we used the 24B GAL4 driver to express UAS-Flipase (UAS-Flp), in the interfollicular stalks (arrows Fig. 4B) to generate clones specifically in these cells. Although this does result in the majority of clones being found in the stalks, it also produces a few clones in the main follicle cells (Fig. 4C). This suggests that either 24B-GAL4 is also expressed at low levels in the follicle cells (which cannot be detected when 24B-GAL4 is used to drive the expression of UAS-LacZ (Fig. 4B)) or that the stalk cells are still capable of becoming follicle cells when they first begin to differentiate. Nevertheless, these clones allowed the evaluation of the role of Mys in the interfollicular stalk precursors.

Interfollicular stalks containing *mys* mutant cells are malformed compared to wild-type stalks of a similar age (Fig. 4C and D). This does not appear to be due to a failure of these cells to differentiate, as Lamin C and 24B-GAL4 are still expressed in mutant cells in the disorganised stalks (Fig. 4E & Fig. S4B). Eya expression is also down-regulated in both mutant and wild type cells (Fig. 4F). Instead of forming single lines of cells, the interfollicular stalks containing *mys* clones are a disorganised mass that can contain wild type as well as mutant cells (Fig. 4G and H). Fully mature wild type stalks are made up of cells with a rectangular cross section, whereas the *mys* mutant cells are round in appearance and usually lie in the centre of the cluster (Fig. 4G-I). Dlg, however, is still only enriched in the regions where cells contact their neighbours (Fig. 4G’&H’), while aPKC expression is absent in the more mature disorganised stalks, as seen in wild-type interfollicular stalks (Fig. 4J). Thus, the intercalation process does not occur correctly in the absence of integrin function, resulting in disordered cell masses several cells thick.

Our observation that Mys is required for the morphogenesis of interfollicular stalks raised the possibility that the areas of epithelial disorganisation at the ends of each egg chamber are in fact misshapen stalks. To test this hypothesis, we generated randomly positioned *mys* clones (using a heat shock-Flp), while also driving lacZ expression with the stalk-specific driver 24B-GAL4. The cells in the disorganised regions of the follicular epithelium at the termini did not express 24B-GAL4, indicating that they are not formed by malformed interfollicular stalks, which abut them posteriorly (Fig. 4K).

When younger egg chambers were examined (before interfollicular stalks have fully formed), the region of epithelial disorganisation was only partially composed of 24B-GAL4 expressing cells, which lie posteriorly (Fig. 4L). Thus, *mys* mutants disrupt the morphogenesis of the terminal follicle cells as well as that of the interfollicular stalks. Our results also suggest that areas of disorganisation anterior to the interfollicular stalk are formed from the follicle cells, like those in Fig. 4L (dashed line), that are neither in contact with the germline (to their anterior) nor destined to become the stalk.

### Mys controls somatic cell positioning in the germarium

Because both the *mys* RNAi and early mutant clones demonstrate that the *mys* phenotype arises in the germarium, we analysed the mutant phenotype at these stages in more detail. In wild type germaria, one surface of every somatic cell contacts the ECM, whether they are follicle cell precursors or future stalk and polar cells (Fig. 5A). By contrast, *mys* mutant cells often lose contact with the ECM and become internalised within the germarium (Fig. 5B; white stars). This internalisation means that cells are mispositioned when egg chamber morphogenesis begins, and this is likely to be the cause of the disorganised interfollicular stalks and follicular epithelium at the termini later in development (Fig. 5C).

**Figure 5.**
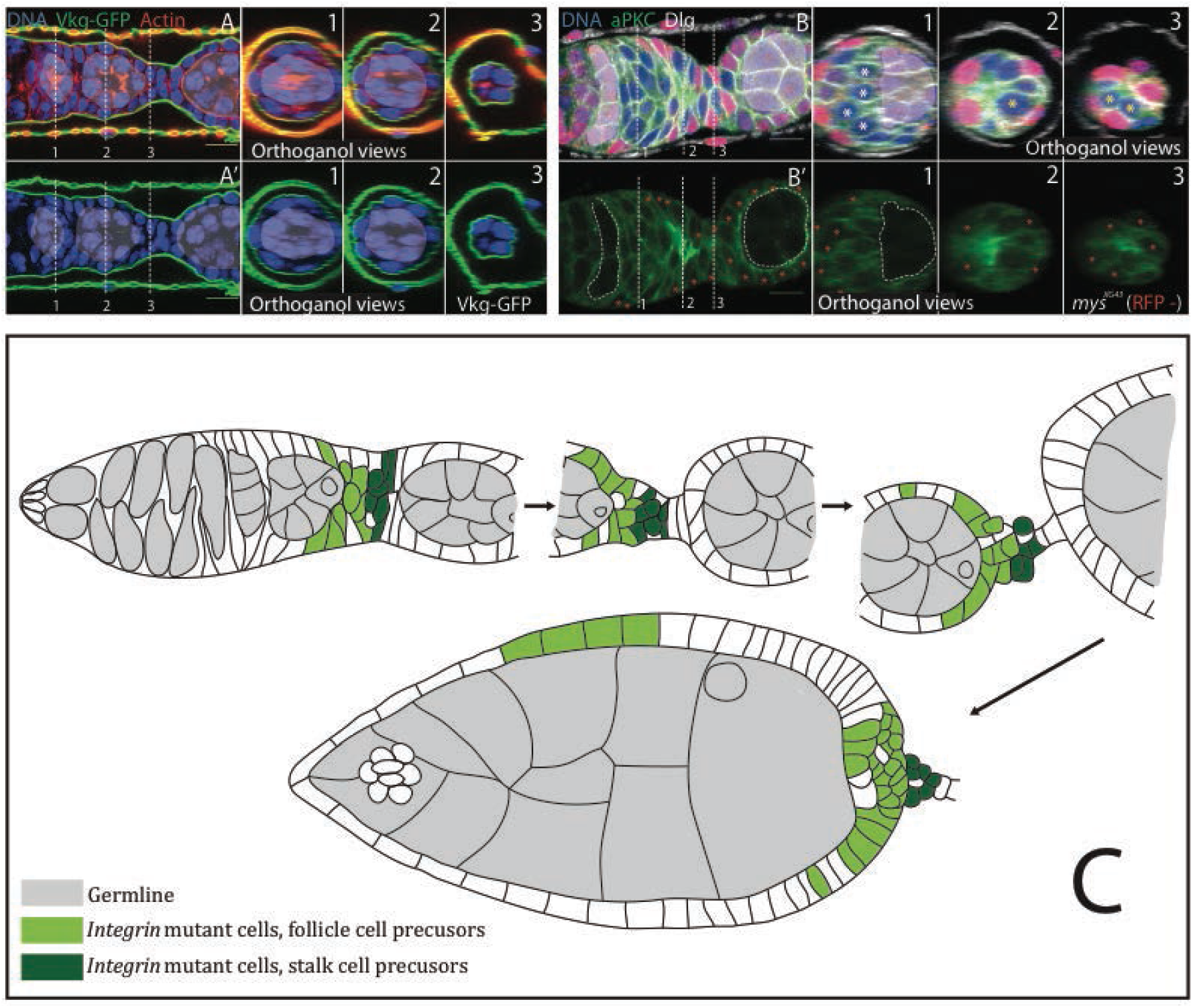
**(A)** The posterior region of a wild-type germarium expressing Viking-GFP (green) to mark the basement membrane and stained for DNA (blue) and F-actin (red). The germline cysts are highlighted by blue shading. The pre-follicle cells form a monolayer of cells that contact the ECM throughout the germarium. The dashed white lines indicate the positions of the orthogonal views shown in the numbered panels to the right. These were generated from Z-stack reconstructions of images taken 0.5μm apart. **(B)** The posterior region of a germarium containing *mys*^*XG43*^ mutant cells marked by the loss of RFP (red and red stars), stained for aPKC (green), DNA (blue in B) and Dlg (white in B). The mutant cells lose contact with the basement membrane and become internalised (white asterix (B)). The dashed white lines indicate the positions of the orthogonal views shown in the numbered panels to the right. These were generated from Z-stack reconstructions of images taken 0.5μm apart. **(C)** A diagram showing the steps in the epithelial disorganisation caused by *mys*^*XG43*^ mutant cells (green) in the FCE. Pale green cells represent those that will end up either in contact with the germline or in disorganised regions at the egg chamber termini. The dark green cells are those that will end up in disorganised interfollicular stalks. The germline cells are shaded in grey. All scale bars are 10μM.

F-actin and the myosin regulatory light chain Spaghetti Squash (Sqh) are basally enriched in pre-follicle cells (Fig. 6A), in contrast to later stage follicle cells where they are enriched at the apical surface (Supp Fig. 4C and D) (Alégot et al., 2018; Finegan et al., 2019). This suggests that the basal sides of the pre-follicle cells are contractile and therefore under tension. Fig. 6B shows 3 cells at various stages of cell internalization, each with a reduced basal surface. Mutant cells typically show an increased basal enrichment of myosin when compared to the adjacent wild-type cells (Fig. 6B and C). This observation suggests that cells respond to the loss of Integrin-ECM adhesion by increasing basal myosin, resulting in a basal contraction that helps to internalise the mutant cells.

**Figure 6.**
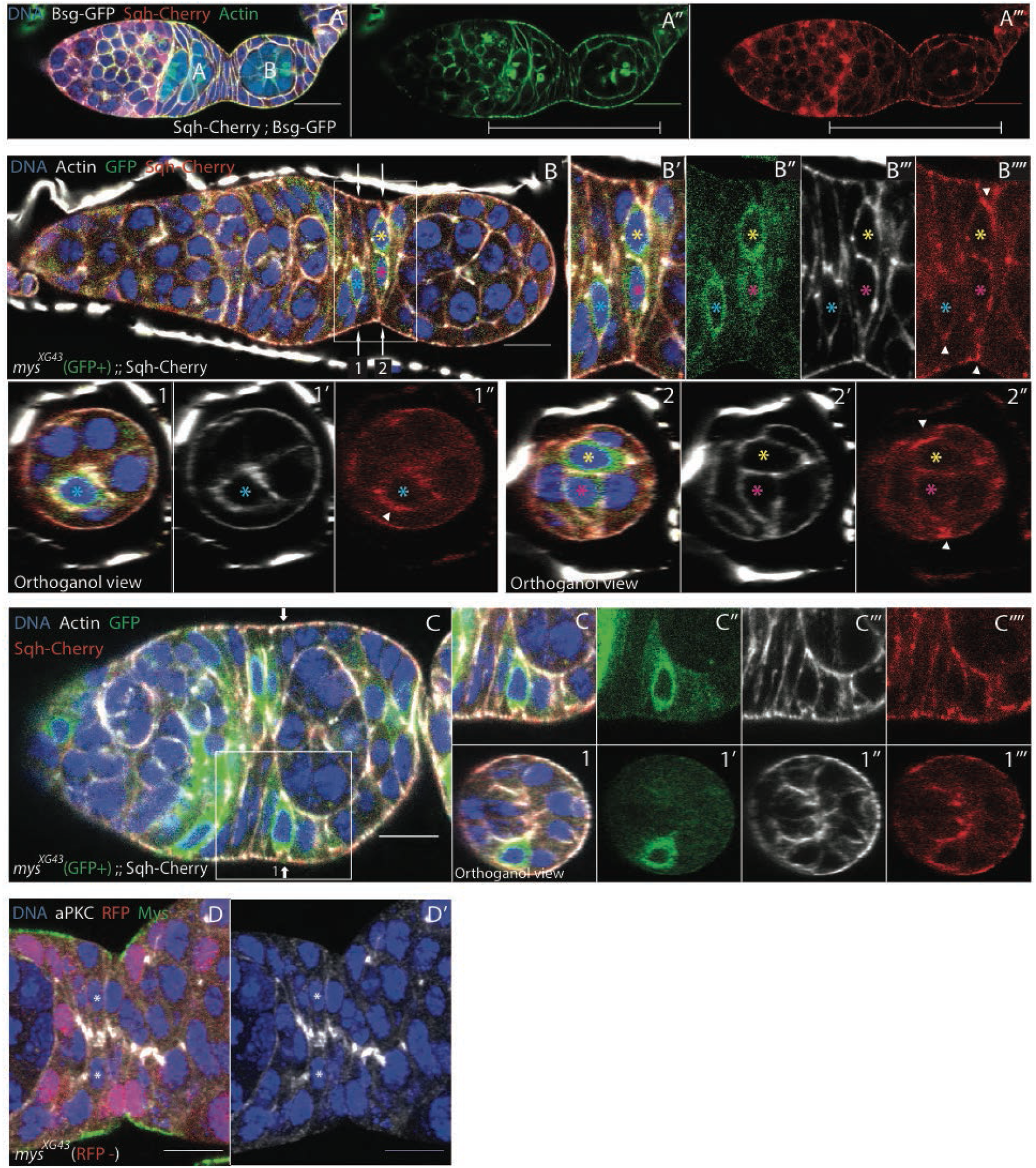
**(A)** A wild-type germarium expressing the myosin regulatory light chain (Sqh) fused to Cherry (red) and Basigin-GFP (white in A), stained for F-actin (red). F-actin and myosin are enriched at the basal sides of the pre-follicle cells (brackets). **(B)** A germarium expressing Sqh-Cherry (red) containing *mys*^*XG43*^ mutant cells marked by GFP (green) and stained for F-actin (white). The mutant cells have detached from the ECM (cells marked with blue, yellow & pink stars). Sqh-Cherry is basally enriched in the detached cells (white arrow heads in B’’’’ and orthogonal views 1’’& 2’’). The positions of the orthogonal views in 1-1’’ and 2-2’’ are indicated by the arrows in B. These are Z stack reconstructions of images taken 0.5μM apart. **(C)** A germarium expressing Sqh-Cherry (red) containing *mys*^*XG43*^ mutant cells marked by GFP (green) and stained for F-actin (white). *mys*^*XG43*^ mutant cells (GFP +) in contact with the germline appear to detach from the ECM. Orthogonal views 1-1”’ are Z stack reconstructions of images taken 0.5μM apart taken at the position of the white arrow in (C). **(D)** A section of an ovariole containing *mys*^*XG43*^ mutant cells marked by the loss of RFP (red) stained for aPKC (white), Mys (green) and DNA (blue). The mutant cells in the stalk region (marked by white asterisks) still localise aPKC to their apical sides. All scale bars are 10μM.

Finally, we asked whether the loss of integrin disrupts the polarity of the interfollicular stalk cells, since these cells never contact the germline and therefore cannot receive the germline-dependent apical polarity cue. We found that *mys* mutant cells still localise aPKC to their presumptive apical sides even if they have detached from the ECM on the opposite side (Fig. 6D). Furthermore, aPKC still localises to the apical sides of mutant cells when two mutant cells are facing each other (Stars Fig 6D). Thus, integrins are also not required for the establishment of apical-basal polarity in the interfollicular stalk cells, despite their lack of an apical cue.

Finally, we asked whether the loss of integrin disrupts the polarity of the interfollicular stalk cells, since these cells never contact the germline and therefore cannot receive the germline-dependent apical polarity cue. We found that *mys* mutant cells still localise aPKC to their presumptive apical sides even if they have detached from the ECM on the opposite side (Fig. 6D). Furthermore, aPKC still localises to the apical sides of mutant cells when two mutant cells are facing each other (Stars Fig 6D). Thus, integrins are also not required for the establishment of apical-basal polarity in the interfollicular stalk cells, despite their lack of an apical cue.

## Discussion

Although it has long been known that integrins are required for the correct organisation of the follicular epithelium at the egg chamber termini, the reason for this defect has remained unclear. Here we show that disorganization in *mys* mutant tissue arises from the failure of the pre-follicle cells to remain attached to the ECM in the germarium. Integrins are not required to maintain the organisation of the epithelium once it has formed. Our analysis reveals that all pre-follicle cells remain in contact with the ECM of the basement membrane as they surround a new germline cyst to form an egg chamber. Loss of Integrin function results in mutant cells that delaminate from the ECM and become internalised, which disrupts the arrangement of cells in a single layer (Fig. 7). As development continues, cells that contact a germline cyst are able to develop as normal, as are any mutant cells that are generated once an egg chamber has been formed. However, mutant cells that lie between germline cysts cannot undergo the normal morphogenetic movements. Signalling from the posterior germline cyst to induce interfollicular stalk cell fate occurs normally despite the altered arrangement of cells, but these cells do not rearrange to form a single-cell wide stalk. Finally, those disordered regions that neither become stalk nor are in contact the germ line develop into the disorganised regions of cells seen at the termini of later-stage egg chambers.

**Figure 7.**
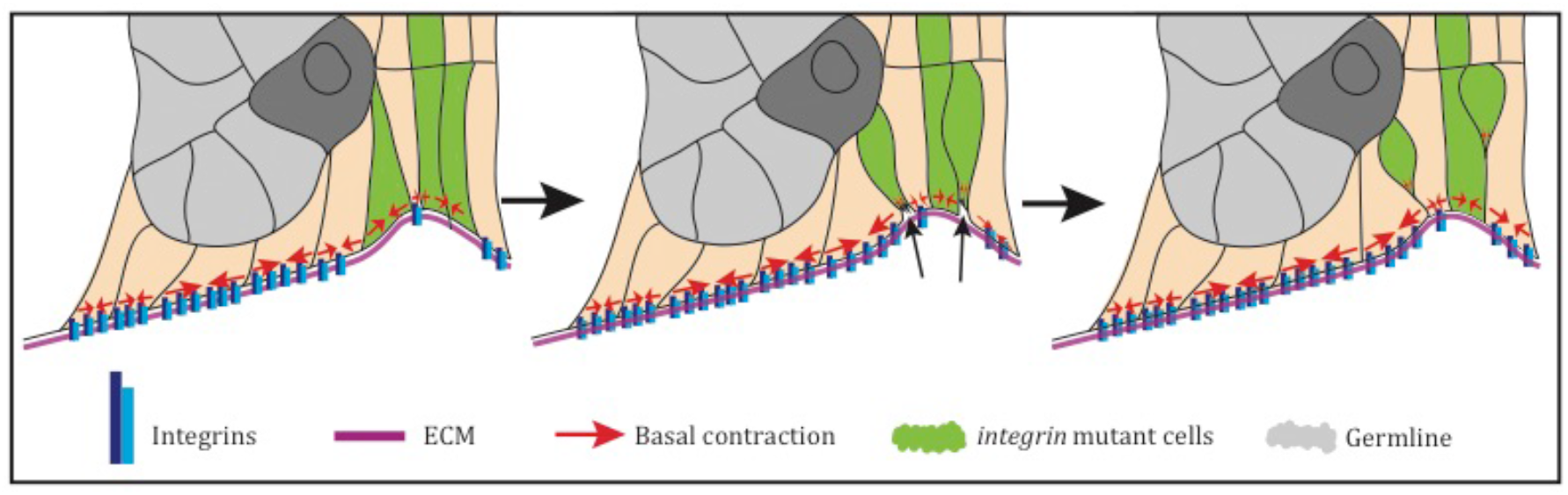
A diagram showing how *mys* mutant cells become internalised. The epithelial disorganisation seems to result from a failure of mutant cells (green) to withstand basal contractions, since these cells are not stably connected to the ECM. The cells are therefore unable to maintain their basal area and undergo a basal constriction (arrows middle panel), causing them to be drawn into the tissue.

Those mutant pre-follicle cells that lose their connection to the ECM appear to undergo a basal constriction, leading to them delaminating from the monolayer into the interior of the tissue. This is similar to a proposed cell extrusion process, in which a cell that becomes detached from the ECM is forced out of the tissue by its neighbours, which spread to fill the gap (McCaffrey and Macara, 2011). However due to the architecture of this particular tissue, the “extruded cells” move into the tissue rather than out of it. As the mutant pre-follicle cells delaminate, the basal constriction is often associated with increased levels of basal myosin, suggesting that increased myosin contractility facilitates the invagination of the mutant cells. This raises the possibility that integrin adhesion to the ECM normally represses basal actomyosin contractility, as has been observed in tissue culture cells (Arthur and Burridge, 2001; Arthur et al., 2000). In support of this view, increasing the levels of the myosin regulatory light chain, Sqh, significantly enhances the epithelial misorganisation phenotype of *mys* mutant clones at the termini (Ng et al., 2016).

The integrin adhesion complex is required for apical-basal polarisation of the other secondary epithelium in *Drosophila*, the midgut, as well as a number of vertebrate epithelia, where it is thought to transduce a basal polarity cue from the basement membrane (Tepass and Hartenstein, 1994; Yu et al., 2005; Akhtar and Streuli, 2012; Chen and Krasnow, 2012;) Chen et al., 2018;). By contrast, integrins appear to perform a purely adhesive function in the pre-follicle cells and play no discernable role in their polarisation. The follicle cells that contact the germ line polarise in response to an unknown apical cue (Tanentzapf et al., 2000).

However, *mys* mutant follicle cells that fail to contact the germ line still manage to partially polarise by generating distinct basal and lateral domains, as revealed by Dlg localisation. Furthermore, a distinct basal domain still forms when the second major ECM receptor, Dystroglycan, is also removed. Thus, integrins and Dystroglycan do not function as redundant receptors for a basal polarity cue from the ECM. One possibility is that there is another ECM receptor that transduces the basal polarity signal. Alternatively, Dlg may be targeted to lateral membranes in response to cell-cell adhesion. In this scenario, there would be no basal polarity cue and instead basal would be specified by the absence of cell-cell contact.

It is even more surprising that the interfollicular stalk precursors polarise normally in the absence of integrins, since these cells never contact the germ line and therefore cannot receive an apical polarity cue, yet they still localise aPKC to the apical domain. This suggests that the cells read some basal cue that generates a polarity that targets apical factors, such as aPKC, to the opposite side of the cell, but the underlying mechanisms are entirely unknown.

The formation of the stalk provides an interesting example of how apical-basal polarity can be remodelled during morphogenesis. The stalk forms from a tube that is initially two cells wide, in which each cell forms an apical domain facing the cell on opposite side. As the cells move past each other to form a one-cell diameter rod, aPKC and E-cadherin do not remain at the leading “ apical” edge of each cell, but instead form a lateral spot that remains approximately in the centre of the stalk, before they eventually disappear. Stalk formation is essentially a process of cell intercalation that resembles the initial steps in the formation of the notochord. In Ascidians for example, presumptive notochord cells intercalate to transform a two-cell wide rod into a line of single cells (Michael T Veeman, 2016; Oda-Ishii et al., 2010). Furthermore, integrin-mediated adhesion to the basement membrane plays a key role in both stalk and notochord morphogenesis (Buisson et al., 2014; Pulina et al., 2014). The stalk could therefore provide a genetically tractable model for investigating how such cell intercalation processes occur.

## Materials and Methods

### Mutant alleles

The following *Drosophila melanogaster* mutant alleles were used: *mys*^*xG43*^ {Wieschaus:1984cd} *dg*^*086*^{Christoforou:2008db}, recombined with FRT19A and FRTG13 respectively. Both stocks were combined to allow the generation of clones mutant for both *mys* and *dg* in the same egg chamber.

### Fluorescent stocks

Mys-GFP and Talin-GFP were gifts from the Brown lab {Klapholz:2015je}. Viking-GFP was produced by the Flytrap project (G00454II) {Morin:2001er}. Mew-YFP (CPTI 001678) and Bsg-GFP (CPTI 100005) were produced by the Cambridge Protein Trap Insertion project (Lowe et al., 2014). FasII-GFP (FasII^397^) is an exon trap insertion line {Silies:2010fd}. Sqh-mCherry is a transgene with a native promoter described in {AdamCMartin:2009du}. E-Cadherin-GFP {Huang:2009ei} and GFP-aPKC {Chen:2018kw} are tagged versions of the endogenous proteins generated by homologous recombination.

### GAL4 drivers

UAS-RNAi constructs against Mys were expressed under the control of GR1-Gal4 {Tran:2003jq} and Traffic Jam-Gal4 {Pancratov:2013eg}. 24B-Gal4 {Brand:1993vw} was used to drive constructs (UAS-LacZ and UAS-mCD8::GFP) specifically in the interfollicular stalks.

### UAS lines

The UAS-Mys-shRNA lines TRiP.JF02819 (Bloomington 27735) and TRiP.HMS00043 (Bloomington 33642) were used to deplete Mys and were generated by the Transgenic RNAi Research Project {Perkins:2015ex}. UAS-LacZ and UAS-mCD8::GFP was used to show expression patterns of the above Gal4 lines.

### FRT lines

RFPnls, hs-FLP, FRT19A was used to generate negatively marked (RFP minus) *mys* mutant clones (Bloomington 31418). w, FRT19A, Tub-Gal80, hsFLP; UAS-lacZ, UAS-mCD8::GFP/CyO; Tub:Gal4/ TM6B (produced in the St Johnston lab) was used to generate positively marked (GFP) *mys* mutant clones. hs-FLP; FTRG13 GFP was used to generate negatively mark (GFP minus) *dg* mutant clones. hs-FLP; FTRG13 GFP and RFPnls, hs-FLP, FRT19A were combined to allow mutant clones of both *mys* and *dg* to be produced simultaneously.

### Other Stocks

*w*^*1118*^ (Bloomington 5905 and 6326) and *y,w* (Bloomington 1495) were both obtained from Bloomington.

### Reagents

The following antibodies were used in this study: mouse anti-Fasciclin II (1D4), and mouse anti-Discs large (4F3), rabbit anti-Myospheroid (CF.6G11), mouse anti-Lamin C (LC28.26), mouse anti-Eyes absent (eya10HB), mouse anti-Armadillo (N2 7A1) (all Developmental Studies Hybridoma Bank), rabbit anti-aPKC (Sigma P0713), rabbit anti-β Galactosidase (MP Biomedicals 0855976), rabbit anti-Lethal giant larvae (Santa Cruz sc-98260) and mouse anti-Fasciclin III (gift from C. Goodman) Rhodamine-Phalloidin, FITC-Phalloidin and 647-Phalloidin were purchased from Invitrogen. Vectashield with DAPI was purchased from Vector Labs. Conjugated secondary antibodies were purchased from Jackson Immunoresearch.

### Immunostaining

Ovaries were fixed for 15 minutes in 4% Paraformaldehyde and 0.2% Tween in Phosphate Buffered Saline (PBS). Ovaries were then incubated in 10% Bovine Serum Albumin (in PBS with 0.2% Tween) to block for at least one hour at room temperature. Primary and secondary immunostainings lasted at least 3 hours in PBS with 0.2% Tween. Three washes (approximately 10 minutes each) in PBS-0.2% Tween were carried out between stainings and after the secondary staining. Primary antibodies were diluted 1:200. Secondary antibodies and Phalloidin were diluted 1:200.

### Imaging

Imaging was performed using an Olympus IX81 (40×/1.3 UPlan FLN Oil or 60×/1.35 UPlanSApo Oil). Images were collected with Olympus Fluoview Ver 3.1. with the exception of Fig. S2H, which was imaged using a Leica MZ16F dissecting microscope fitted with a Qimaging Qiclick camera and collected using QCapture Pro 6.0. Image processing was preformed using Image J.

### *Drosophila* genetics

Follicle cell clones of *mys*^*XG43*^ and *dg*^*086*^ were induced indiscriminately both alone and simultaneously by incubating hs-FLP; FRT mutant/ FRT marker larvae or pupae at 37°C for two out of every twelve hours over a period of at least two days. Apart from the heat shocks, the crosses were raised at 25°C. Adult females were dissected at least two days after the last heat shock. *mys*^*XG43*^ clone production was targeted to the interfollicular stalk cells by using 24B-Gal4 to drive the expression of UAS-FLP. Crosses were raised at 25°C and adult females were dissected at least two days after they had eclosed. RNAi lines against Mys were crossed with TJ-Gal4 and either raised solely at 25°C (Fig 2E and F, Fig 3B, Fig S2C and H), solely at 18°C (Fig S2D and E) or at 18°C and then adult flies were shifted to 29°C for 2 days before dissection (Fig 2C, D, Fig S2F & G). RNAi lines against Mys were crossed with GR1-Gal4 and had to be raised at 18°C in order to produce adults, which were then moved to 29°C for 2 (Fig 2C & D) or 6 days (Fig S2A) before dissection.

## Acknowledgements

We thank Nick Brown and his lab for fly stocks and Corey Goodman for the FasIII antibody; the Developmental Studies Hybridoma Bank, Kyoto Stock Center (DGRC) and Bloomington Drosophila Stock Center (BDSC) for antibodies and fly stocks; The Piddini and Brown labs for helpful discussion and the D. St J. lab, in particular Nick Lowe and Jackie Hall, for technical assistance, helpful comments and criticism.

## Author contributions

H. E. L., D. T.B. and D.St J. conceived the project and designed the experiments. H.E.L. performed the experiments. H. E. L and D. St J. wrote the manuscript.

## Supplementary Figures

**Figure S1.**
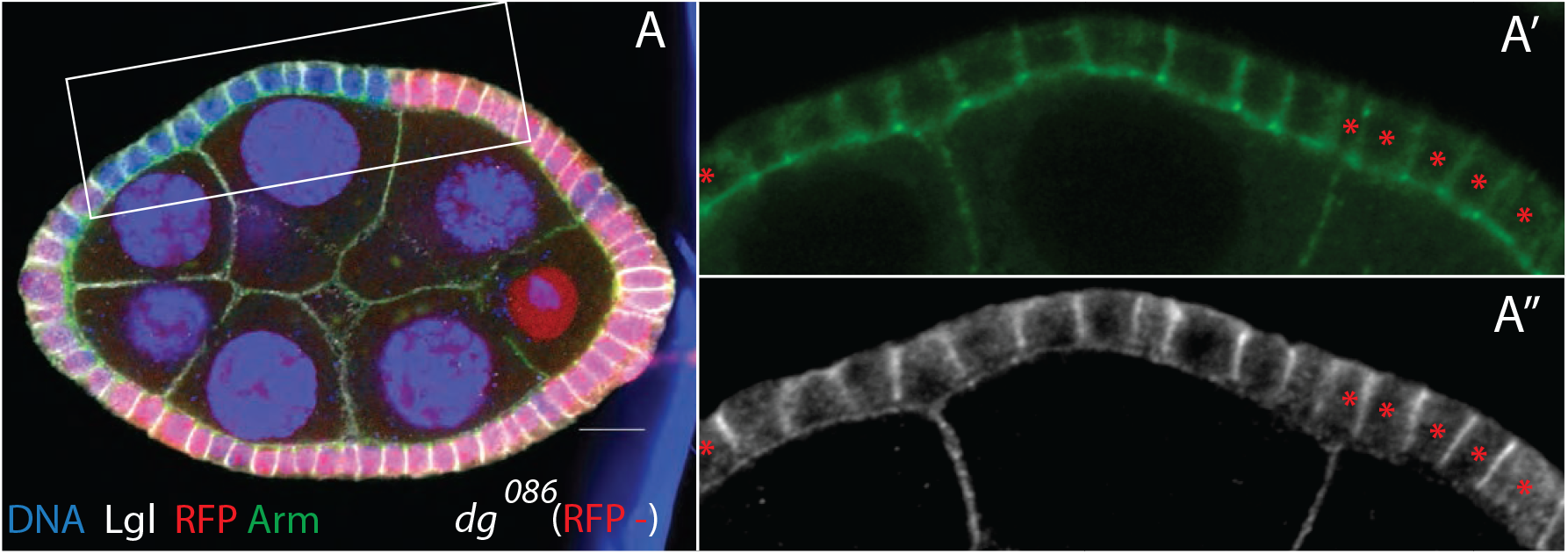
**(A)** A stage 7 egg chamber containing *dg*^*086*^ mutant cells marked by loss of RFP (red) and stained for Armadillo (green) and Lgl (white). The mutant cells show neither epithelial disorganisation nor a disruption in apical-basal polarity when they occur along the sides of egg chambers. Scale bar is 10μM.

**Figure S2.**
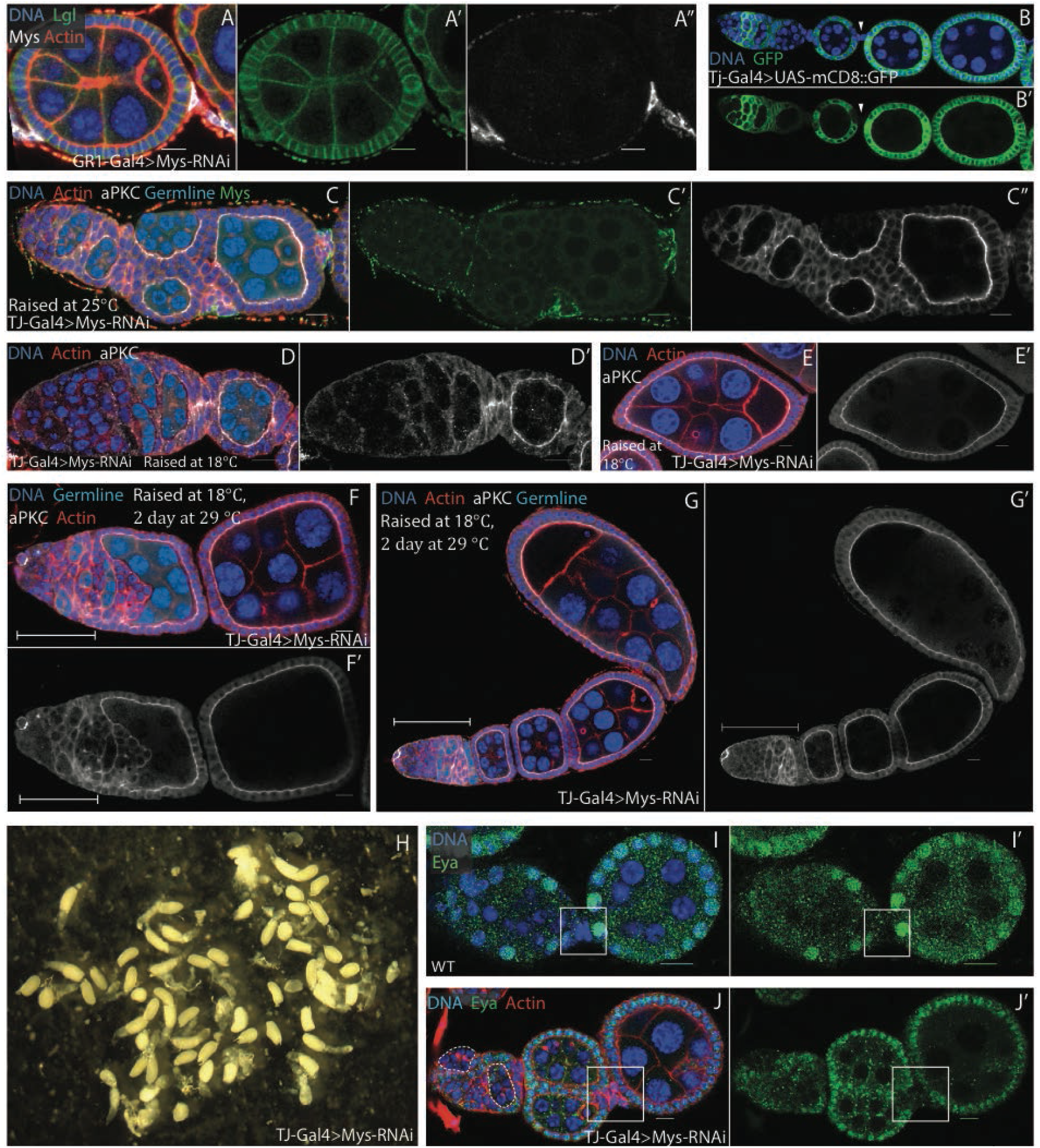
**(A)** A stage 4 egg chamber expressing UAS-*mys*-RNAi under the control of GR1-Gal4 stained for Mys (white), Lgl (green), F-actin (white) and DNA (blue). The flies were raised at 18°C and moved as adults to 29°C for 6 days before dissection. Although Mys is efficiently depleted from the follicle cells, this does not result in a loss of tissue organisation or apical-basal polarity. **(B)** A germarium expressing mCD8-GFP (green) under the control of TJ-Gal4, which drives the expression of UAS constructs in all somatic cells of the ovary, apart from the interfollicular stalks (white arrow heads). **(C)** A germarium from flies raised at 25°C expressing UAS-*mys*-RNAi (Bloom 27735) under the control of TJ-Gal4, stained for Mys (green), aPKC (white), F-actin (red) and DNA (blue). Mys is efficiently knocked down in all somatic cells (except interfollicular stalk cells), resulting in gross disorganisation of the ovariole. The follicle cells that contact the germ line polarise correctly as shown by the apical localisation of aPKC. **(D and E)** Germaria from TJ-Gal4; UAS-*mys*-RNAi (Bloom 27735) flies raised at 18°C. At this temperature, TJ-GAL4 does not drive sufficient *mys*-RNAi expression to produce a phenotype, resulting in normal germaria (D) and egg chambers (E). **(F and G)** Germaria from TJ-Gal4; UAS-*mys*-RNAi (Bloom 27735) flies raised at 18°C and then shifted to 29°C for two days before dissection, aPKC (white), F-actin (red) and DNA (blue). The early germine cysts are outlined in pale blue. Knock-down of Mys disrupts the organisation of the follicle cells in the germarium (brackets F and G) but does not affect older egg chambers. **(H)** Eggs from TJ-Gal4; UAS-*mys*-RNAi (Bloom 27735) flies raised at 25°C. Many of the eggs are misshapen and rounder than normal, consistent with a defect in egg chamber rotation. **(I)** Eya staining in wild-type. Eya is expressed in the main body follicle cells, but not the polar or stalk cells (box). **(J)** An ovariole expressing UAS-*mys*-RNAi (Bloom 33647) under the control of TJ-Gal4 at 25°C, stained for Eya (green), F-actin (red) and DNA (blue). Eya negative cells can be found between developing germline cysts (box), indicating that stalk and polar cells have been specified normally. Early germline cysts are outlined with dashed lines. All scale bars are 10μM.

**Figure S3.**
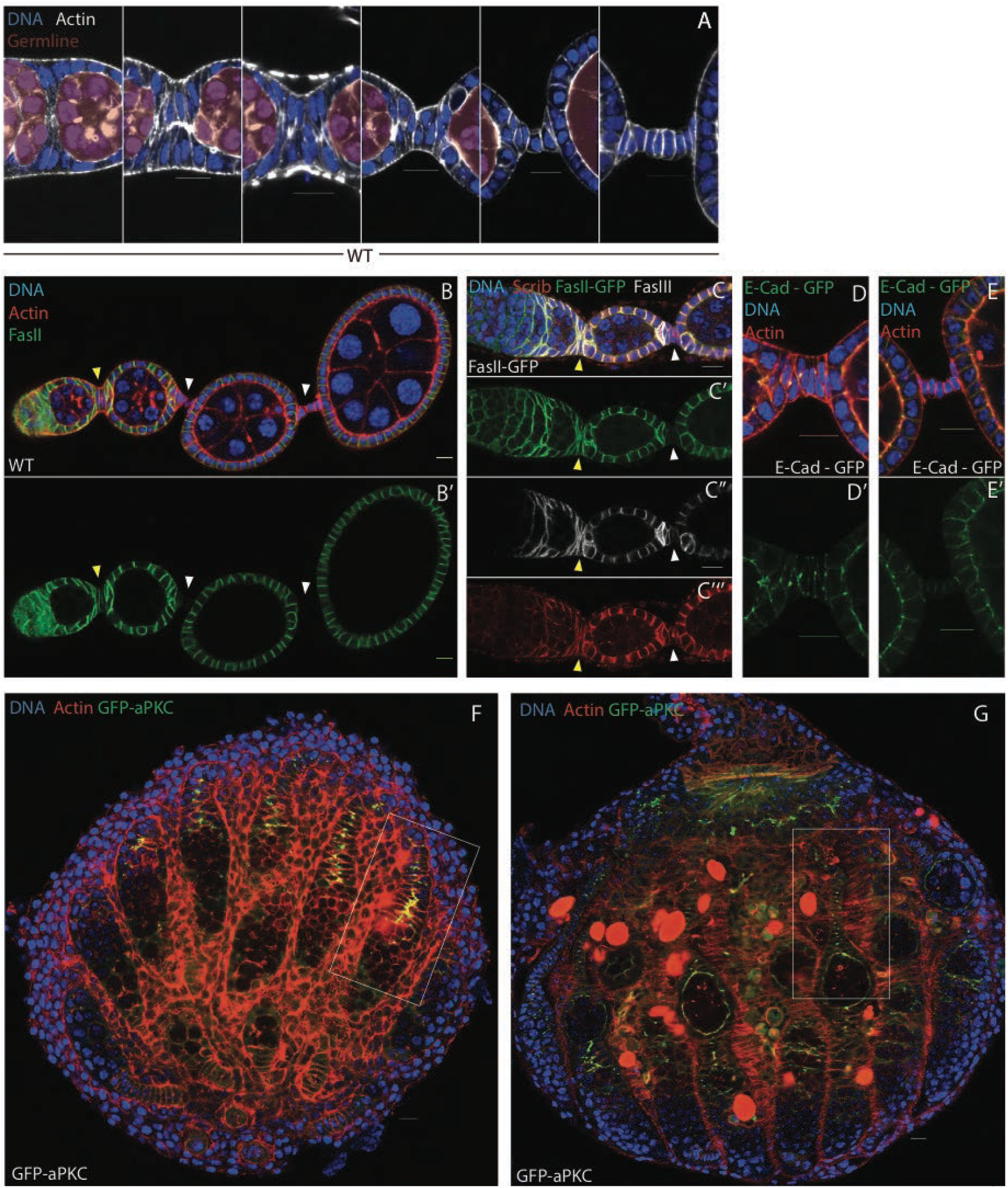
**(A)** Series of interfollicular stalks of increasing age (left to right) stained for F-actin (white) and DNA (blue). Interfollicular stalks are made from two rows of cells between adjacent germline cysts that intercalate to form a one cell wide stalk. **(B)** A wild-type ovariole stained for FasII (green), F-actin (red) and DNA (blue). FasII localises laterally domain in follicle cells and the developing stalk (yellow arrowhead), but is down regulated later as the stalks mature (white arrowheads). **(C)** A wild-type ovariole expressing FasII-GFP (green) and stained for FasIII (white), Scribbled (red) and DNA (blue). FasIII is down-regulated as interfollicular stalks mature (yellow vs white arrow head), but at a later time point than FasII (compare white arrowheads in C’ & C’’). **(D and E)** Developing stalks expressing E-Cad-GFP (green) and stained for actin (red) and DNA (blue). Like aPKC, E-Cad localises to lateral punctae as the stalk cells intercalate (D) and is down-regulated as the stalk matures (E). **(F)** A developing pupal ovary expressing aPKC-GFP (green) and stained for actin (red) and DNA (blue). aPKC localises to the apical sides of the basal stalk cells prior to intercalation. Enlarged image of box is in Fig. 3I **(G)** An older pupal ovary expressing aPKC-GFP (green) and stained for F-actin (red) and DNA (blue), aPKC relocalises to lateral punctae as the basal stalk cells intercalate. Enlarged image of box is in Fig. 3J. All scale bars are 10μM.

**Figure S4.**
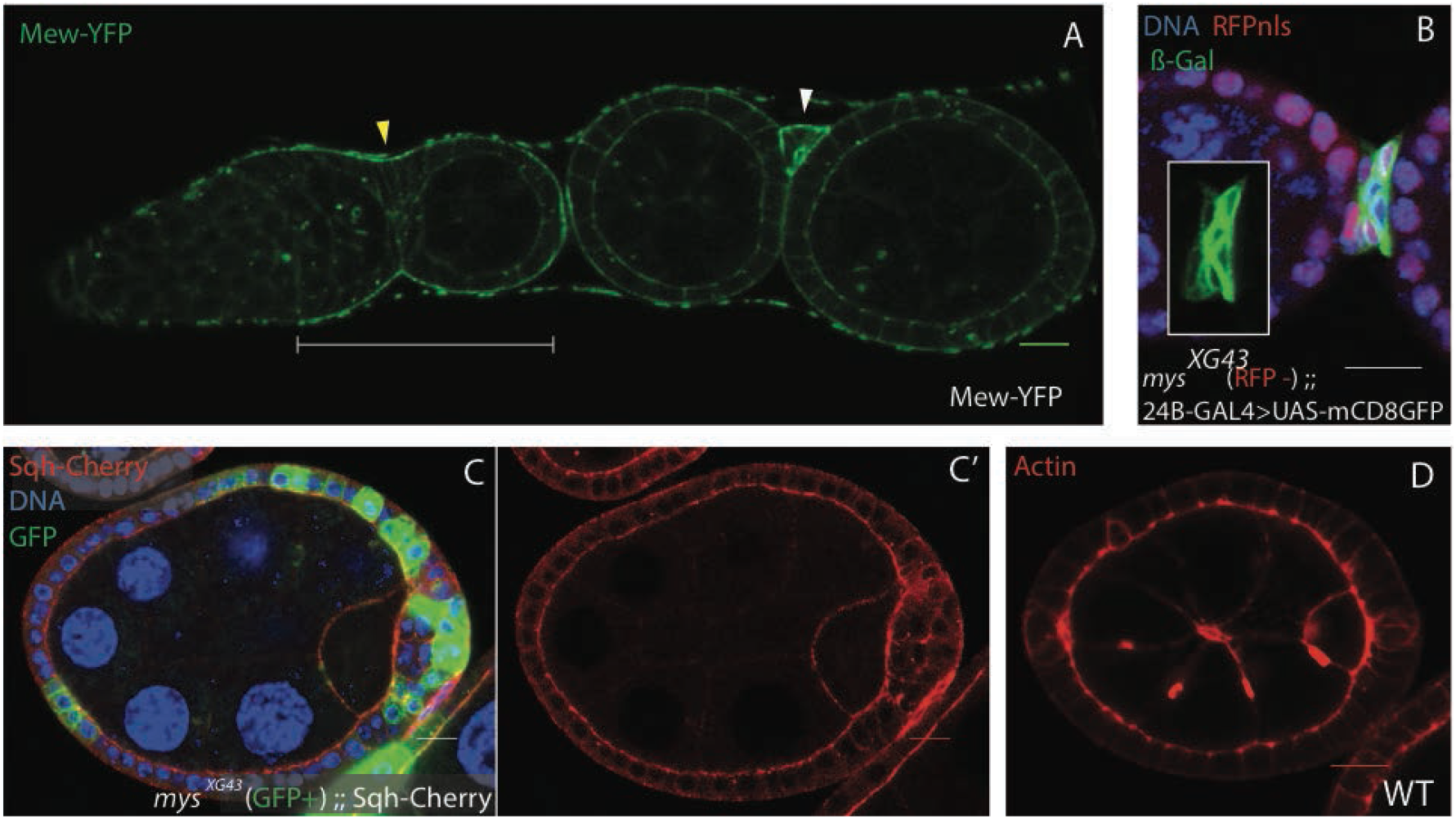
**(A)** An ovariole expressing Mew-YFP (green) from a BAC genomic construct. Mew is basally enriched in the germarium (bracket) and in the forming (yellow arrow head) and fully produced (white arrow head) interfollicular stalks. **(B)** The stalk region of an ovariole containing *mys*^*XG43*^ mutant cells marked by the loss of RFP (red) and expressing UAS-LacZ (β-galactosidase, green) and UAS-Flp under the control of the stalk driver 24B-Gal4. Stained for DNA (blue). Although the morphogenesis of the stalk is disrupted, the stalk marker 24B-Gal4 is expressed in *mys*^*XG43*^ mutant and wild type cells. **(C)** An egg chamber containing *mys*^*XG43*^ mutant cells marked by the expression of GFP (green) and expressing Sqh-Cherry (red) and stained for DNA (blue). Sqh is expressed and localises (apically enriched) normally in *mys*^*XG43*^ mutant cells that contact the germline, as well as being expressed at similar levels in cells which do not contact the germline. **(D)** F-actin is apically enriched in wild-type follicle cells All scale bars are 10μM.

